# AlphaMap: An open-source Python package for the visual annotation of proteomics data with sequence specific knowledge

**DOI:** 10.1101/2021.07.30.454433

**Authors:** Eugenia Voytik, Isabell Bludau, Sander Willems, Fynn Hansen, Andreas-David Brunner, Maximilian T. Strauss, Matthias Mann

## Abstract

**Summary:** Integrating experimental information across proteomic datasets with the wealth of publicly available sequence annotations is a crucial part in many proteomic studies that currently lacks an automated analysis platform. Here we present AlphaMap, a Python package that facilitates the visual exploration of peptide-level proteomics data. Identified peptides and post-translational modifications in proteomic datasets are mapped to their corresponding protein sequence and visualized together with prior knowledge from UniProt and with expected proteolytic cleavage sites. The functionality of AlphaMap can be accessed via an intuitive graphical user interface or – more flexibly – as a Python package that allows its integration into common analysis workflows for data visualization. AlphaMap produces publication-quality illustrations and can easily be customized to address a given research question.

**Availability and implementation:** AlphaMap is implemented in Python and released under an Apache license. The source code and one-click installers are freely available at https://github.com/MannLabs/alphamap.

**Supplementary information:** A detailed user guide for AlphaMap is provided as supplementary data.

## Introduction

Bottom-up mass spectrometry has become the leading technology for identifying and quantifying proteomes [1]–[3]. Since peptides rather than intact proteins are measured, the identity and quantity of individual proteins are inferred from their matching peptides. Impressive technical developments in both full proteome data acquisition [4]–[7] as well as in the analysis of different post-translational modifications (PTMs) [8], [9] continue to increase the achievable sequence coverage in bottom-up proteomics workflows. It is of high interest to investigate which sequence regions of a protein are covered by such proteomics experiments and to inspect where experimentally determined PTMs are located. Visualizing identified peptides and PTMs together with known protein sequence information, such as specific protein domains or previously observed modifications retrieved from UniProt [10], is therefore an important aspect of downstream data exploration. However, the ability to easily integrate and visualize experimental data together with already known sequence annotations is an unmet need in the proteomics community. Although established visualization platforms provide manual visualization of a single experimental sample or dataset at the time [11], there is a lack of tools that support state-of-the-art data analysis software frameworks and that can visualize experimental sequence coverage across multiple samples or datasets in combination with available sequence annotations mined from UniProt, the standard knowledgebase for protein information [10]. To make this wealth of information easily accessible to proteomics researchers, we developed AlphaMap, a Python package that facilitates the visual exploration of peptide-level proteomics data, while additionally integrating protein sequence information based on proteolytic cleavage sites and UniProt annotations. By providing both an intuitive graphical user interface (GUI) as well as the Python package itself, AlphaMap can be employed by end users as well as by other developers that aim to integrate its functionality in their own tools, for example in the development of data visualization platforms.

### The AlphaMap computational framework

In line with other recently developed software tools from our lab [12], [13], we implemented AlphaMap in pure Python because of its clear, easy to understand syntax and the availability of excellent supporting scientific libraries. To read fasta files, we leverage the Pyteomics Python package [14], [15]. Plotly is a well-established plotting library that we use for generating AlphaMap’s sequence visualization [16], allowing flexible customization and great user interactivity. To enable easy access to the AlphaMap functionality with a low barrier of entry, a stand-alone GUI was implemented using the Panel library [17]. AlphaMap can be launched either as a browser-based GUI after simple local installation or as a standard Python module installed via PyPI [18] or directly from its GitHub repository.

In line with the AlphaPept ecosystem, we make the AlphaMap code openly available on GitHub, employing its many supporting features for unit and system testing via GitHub actions. For code development, we adopted the concept of ‘literate programming’ [19], which combines the algorithmic code with readable documentation and testing. Using the nbdev package, the codebase can directly be inspected in well documented Jupyter Notebooks, from which the code is automatically extracted [20]. We envision that these design principles will encourage the broader community to integrate AlphaMap in their own data analysis and visualization workflows with the possibility to easily adopt the code according to specific needs.

### Overview of the AlphaMap workflow

AlphaMap uses peptide-level proteomics data as input. It currently supports the direct import of data processed by MaxQuant [21], Spectronaut [22], DIA-NN [23] and our recently introduced AlphaPept framework [12].

In contrast to Protter [11], users can select multiple independent datasets for co-visualization. These could either have been processed by the same or with different MS analysis tools. It is also possible to select only a single sample, or a subset of samples of a given input file for individual sequence visualization. In addition to the peptide-level data generated from LC-MS analysis, AlphaMap leverages a plethora of manually curated sequence-specific protein level information available from UniProt. Fasta files and UniProt sequence annotations are readily accessible in AlphaMap for the 13 most popular UniProt organisms as well as for SARS-CoV and SARS-CoV-2. Functionality to enable the integration of additional organisms is further available as part of our Python package. Finally, the user can select the different layers of information that should be displayed in the interactive sequence representation, including selected protease cleavage sites and UniProt sequence annotations. Figure 1A shows a schematic overview of the AlphaMap workflow. Detailed instructions for its installation and usage are further provided in the supplementary user guide. In addition to interactive sequence visualization of a user-selected protein, AlphaMap provides individual links to external databases and tools for further sequence evaluation in UniProt [10], PhosphoSitePlus [24], Protter [11], PDB [25] and Peptide Atlas [26].

**Figure 1.**
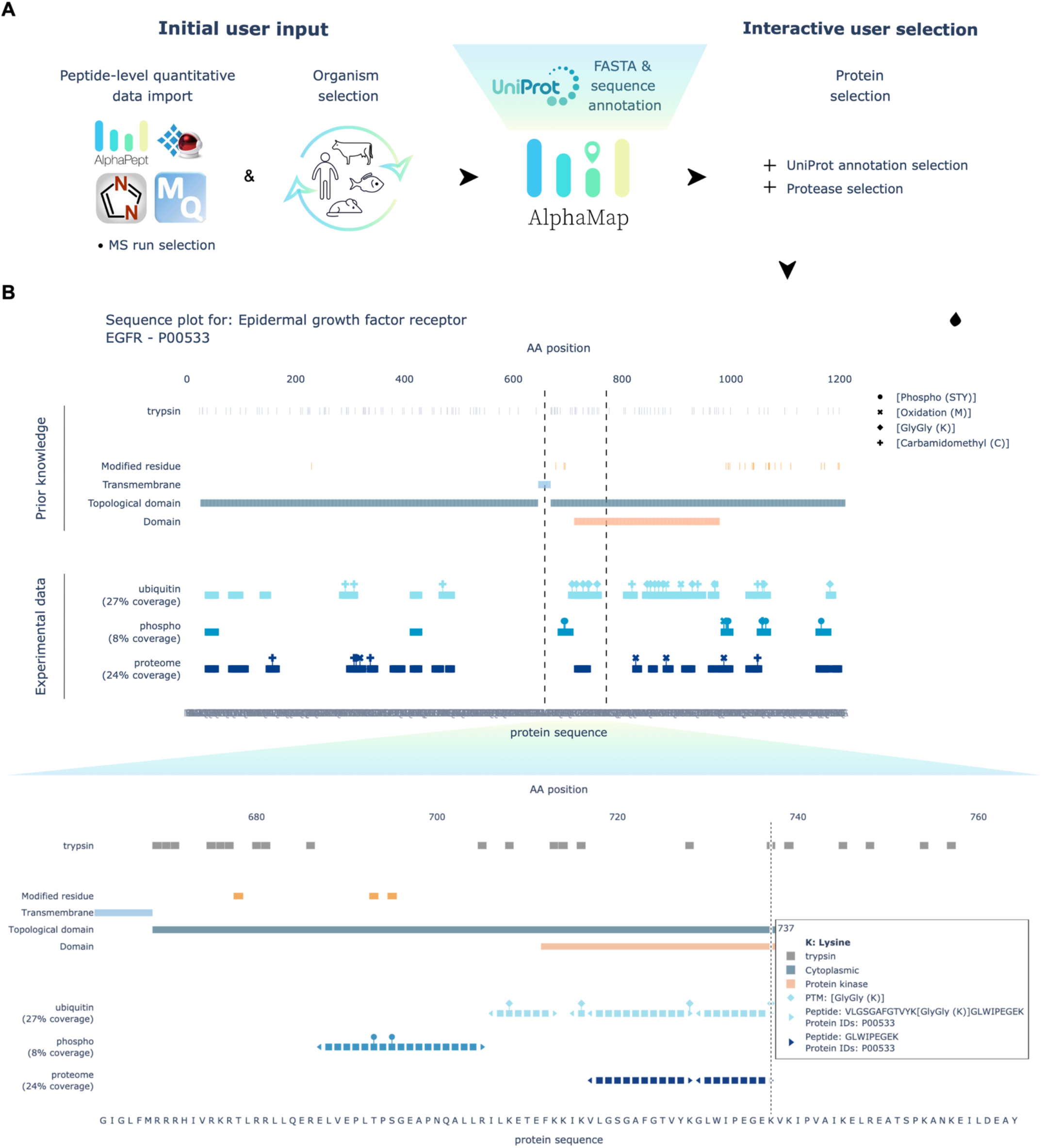
**A** Overview of the AlphaMap workflow from MS data upload to the interactive sequence visualization. **B** Exemplary sequence visualization for epidermal growth factor receptor (EGFR). A zoom-in on a selected sequence region, indicated by dashed lines, is provided at the lower part of the panel.

### Application of AlphaMap to investigate full proteome and PTM data

Figure 1B shows the sequence visualization of the peptides and PTMs identified for the epidermal growth factor receptor (EGFR) in human A549-ACE2 cells that were infected with SARS-CoV-2 or SARS-CoV (an exemplary viral protein detected in this dataset is visualized in the supplement) [27]. We show three independent experimental traces: one for full proteome data, one for phospho-enriched peptides and one for ubiquitin-enriched peptides. The proteome data indicates a homogeneous coverage across the entire protein sequence. As expected, phosphorylation and ubiquitination is limited to the C-terminal region of the protein, which is annotated to be exposed to the cytosol. Additionally, the kinase domain of EGFR is highly ubiquitinated in our dataset, whereas the surrounding cytosolic regions are phosphorylated. Interestingly, AlphaMap reports that most of our observed phosphorylation sites have been previously identified, whereas none of the identified ubiquitination sites are annotated in UniProt.

Beyond the uses highlighted here, we envision AlphaMap to facilitate data analysis and interpretation for a variety of different applications:

- Candidate validation: AlphaMap can be used to assess the sequence coverage of identified biomarker candidates (or other proteins of interest) to evaluate possible sequence variations or unexpected anomalies on the basis of readily available sequence information.
- Preparation of panels for publication: Sequence visualizations form AlphaMap can directly highlight the precise MS derived information about proteins of interest in biological or clinical projects.
- Technical comparisons: AlphaMap can be used to evaluate sequence coverage between different data acquisition strategies such as data-dependent and data-independent acquisition, alternative instrument platforms or software tools.
- Optimization of sample processing: Visualization of protein cleavage sites for different proteases can help to optimize sample processing with the goal to achieve a more complete sequence coverage.

## Conclusion

AlphaMap offers an interactive GUI and a Python package for visualizing peptide-level bottom-up proteomics data on the basis of individual protein sequences, including information of curated UniProt sequence annotations and expected proteolytic cleavage sites. We expect that future developments by us and the community will extend the variety of available annotations in AlphaMap, for example by including prior knowledge of sequence conservation or predicted functional domains. We envision that AlphaMap will assist MS-based proteomics researchers in inspecting peptide- and PTM-level data, thereby providing valuable information in the process of candidate validation in biological and clinical context.

## Supporting information

Supplementary AlphaMap user guide

## Acknowledgements

We thank Julia Schessner, Barbara Steigenberger, Jakob Bader and Sophia Mädler for testing and providing critical feedback on AlphaMap. We are grateful to Özge Karayel and Maria C. Tanzer for valuable discussions and for providing experimental data.

This study was supported by The Max-Planck Society for Advancement of Science and by the Bavarian State Ministry of Health and Care through the research project DigiMed Bayern (www.digimed-bayern.de). IB acknowledges funding support from her Postdoc.Mobility fellowship granted by the Swiss National Science Foundation (P400PB_191046).

## Author contributions

IB conceptualized the project and together with EV and MM wrote the manuscript with contributions from all authors. IB and EV implemented the core AlphaMap functions. EV implemented the GUI. SW provided important help with the AlphaMap installers. FH and ADB provided valuable ideas for the concept and visualization in AlphaMap and FH further contributed by rigorous testing. MTS designed the general AlphaPept ecosystem and assisted with the nbdev environment. MM supervised the study and provided critical feedback on all aspects of the presented work.

