## Supplementary AlphaMap user guide for "AlphaMap: An open-source Python package for the visual annotation of proteomics data with sequence specific knowledge"

### AlphaMap Tutorial

Developed by: Eugenia Voytik and Isabell Bludau

This step-by-step guide helps you to get started with our software AlphaMap.

#### Table of Contents


*................................................................................................ - 1 -*

***Program Description* ........................................................................ - 1 -**

***Installation* ........................................................................................ - 2 -**

**Windows ........................................................................................ - 2 -**

**MacOS ........................................................................................... - 4 -**

***How to use the AlphaMap GUI* .......................................................... - 6 -**


#### Program Description

AlphaMap enables the exploration of proteomic datasets on the peptide level. It is possible to evaluate the sequence coverage of any identified protein and its post-translational modifications (PTMs). AlphaMap further integrates all available UniProt sequence annotations as well as information about proteolytic cleavage sites.

### Installation

#### Windows

1. Download the latest release for Windows (alphamap_installer_windows.exe) from the GitHub repository and open the .exe file.
2. In the "User Account Control" dialog asking about permission for the app to make changes to your device press the "Yes" button.
3. In the appearing "Setup – alphamap version X.X.X" dialog window accept the License Agreement and press the "Next" button.

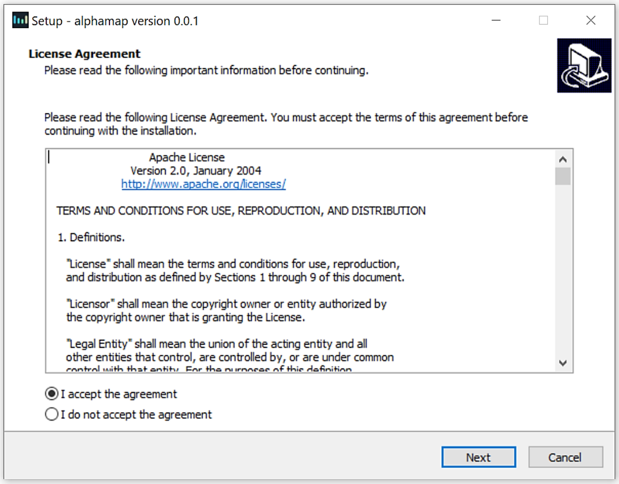


4. Select the destination location for the installation of AlphaMap software (the size of the whole package is 994.6 MB) and press the "Next" button.

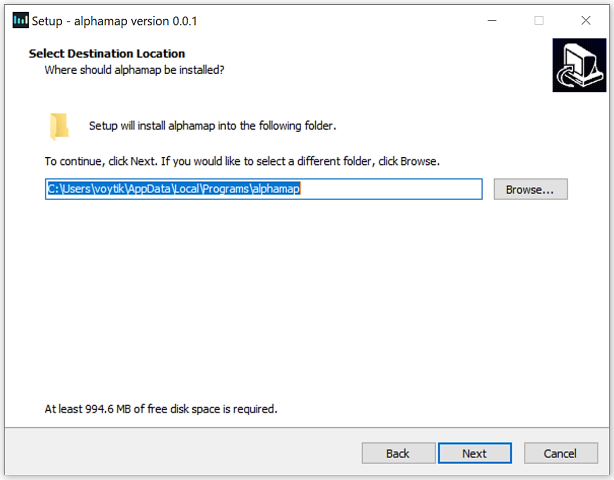

5. In the next dialog window mark the “Create a desktop shortcut” check box and press the “Next” button.

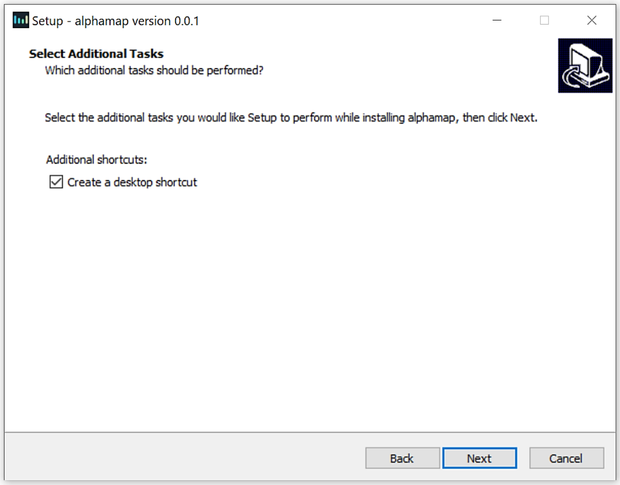


6. Check the setting and if everything is correct, press “Install” button. You may go back to change some settings using the “Back” button or “Cancel” the installation.

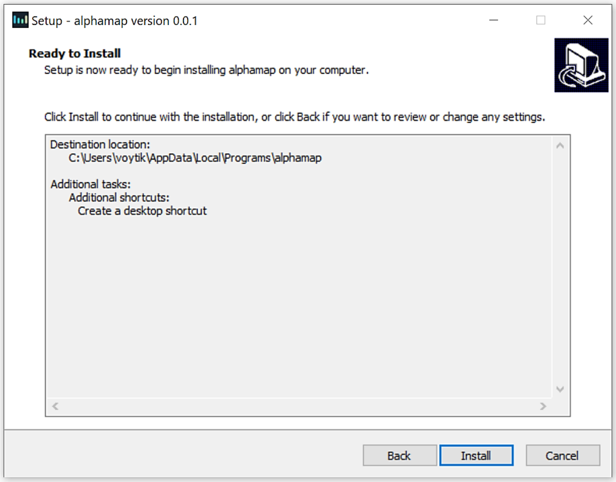


7. Wait till the installation process is finished and with the marked “Launch alphamap” check box press the “Finish” button.
8. In the appearing “Windows Security Alert” dialog window press the “Allow access” button that will prevent the Windows Defender Firewall from blocking the AlphaMap tool on your PC.
9. Check your default browser (Google Chrome or Mozilla Firefox are suggested for the fast running of the AlphaMap) and start working with the tool.

* If you install AlphaMap for all users, you might need admin privileges to run it (right-click on the AlphaMap logo on your desktop and select "Run as administrator").

### MacOS

1. Download [the latest release](#) for macOS (alphamap_gui_installer_macos.pkg) from the GitHub repository and open the .pkg file.
2. Click “Continue” on the appearing “Install AlphaMap X.X.X” dialog window.

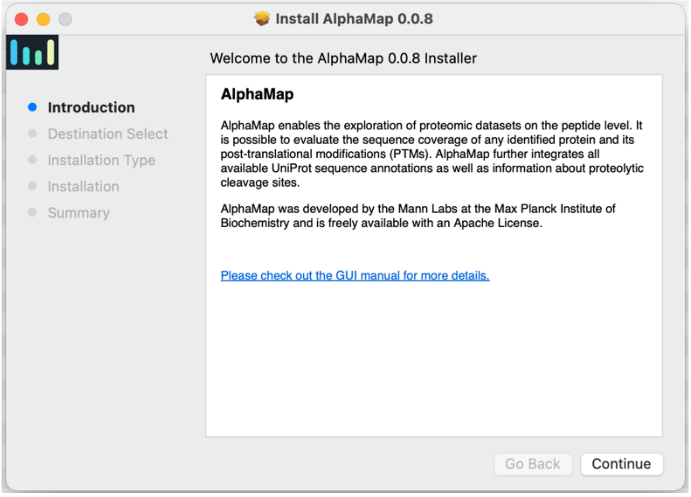


3. Click “Install” to start the installation. This might take a few minutes.

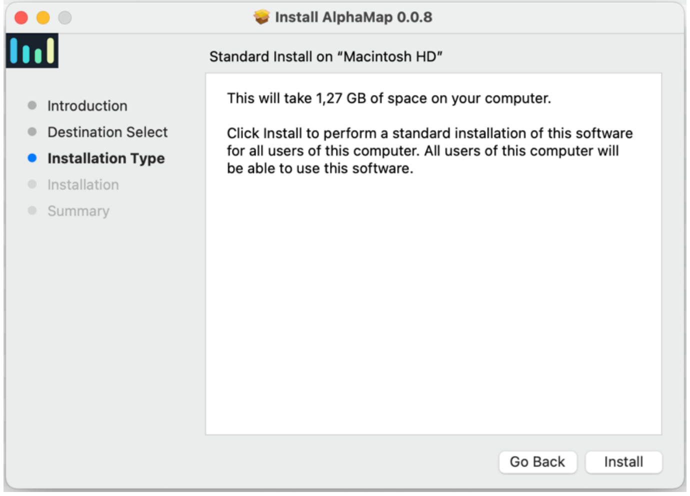


4. Click “Close” to quit the installation menu. AlphaMap is now available in your applications folder on your MacOS.

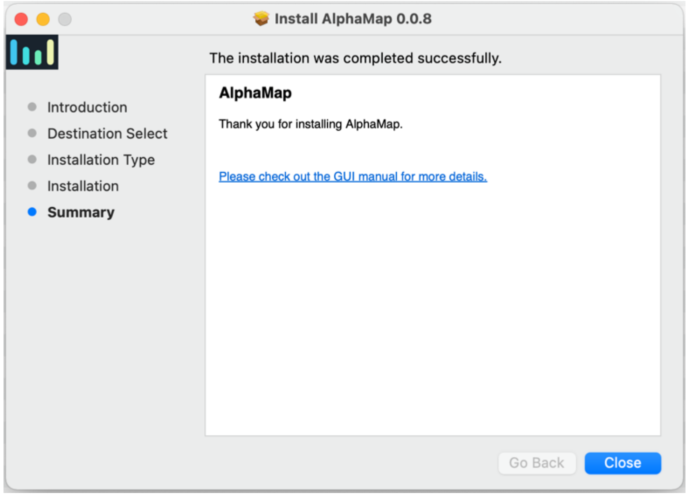


* In rare cases, you might get an error message that the “alphamap_gui_install_macos.pkg” cannot be opened because it is coming from an unidentified developer. This error can be avoided by pressing the OPTION key.

### How to use the AlphaMap GUI

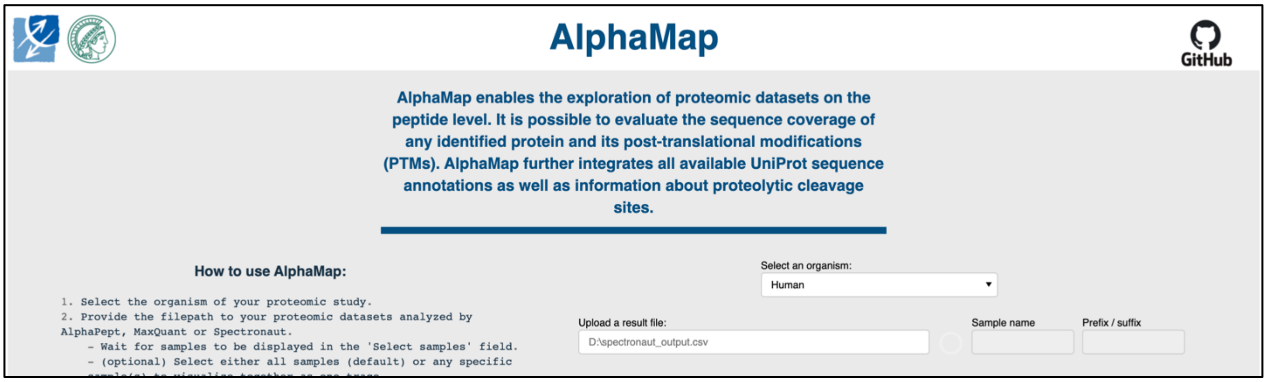


1. First select the organism of your proteomic study. Currently, the [13 most popular organisms based on UniProt] are available for selection, including: Human [Taxon identifier=9606], Mouse [10090], Rat [10116], Cow [9913], Zebrafish [7955], Drosophila [7227], Caenorhabditis elegans [6239], Slime mold [44689], Arabidopsis thaliana [3702], Rice [39947], Escherichia coli (strain K12) [83333], Bacillus subtilis (strain 168) [224308], Saccharomyces cerevisiae (strain ATCC 204508 / S288c) [559292]. Additionally, the reviewed proteomes of SARS-CoV [694009] and SARS-CoV2 [2697049] are available.

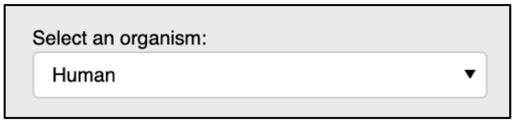


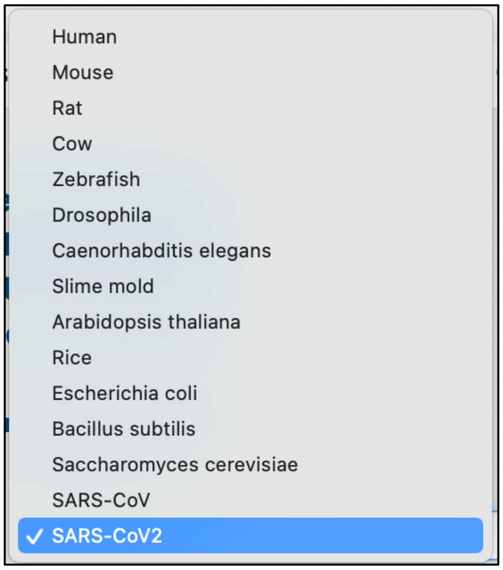

2. Upload your proteomic datasets analyzed by AlphaPept, MaxQuant, Spectronaut or DIA-NN:

    a) Provide the filepath to the result file in the "Upload a result file:" field, e.g. "D:\alphapept_output.csv" (Windows) or "/Users/test/alphapept_output.csv" (MacOS).

    b) Wait for samples to be displayed in the "Select samples" field. The loading process is indicated by a spinner symbol.

    c) (optional) Select either all samples (default) or any specific sample(s) to visualize together as one trace.

    d) (optional) Choose a name by which the selected sample(s) will be displayed in the figure. If no name is provided, the original names of all selected samples will be concatenated by semicolon. If 'all samples' were selected, the filename will be the default name.

    e) (optional) Provide a prefix or suffix to be removed from the original names of the selected samples. This option only applies if no user defined name is provided (see d).

    f) Up to three datasets or sets of selected samples can be visualized together in the GUI. For this, use the "Upload additional result files" option.

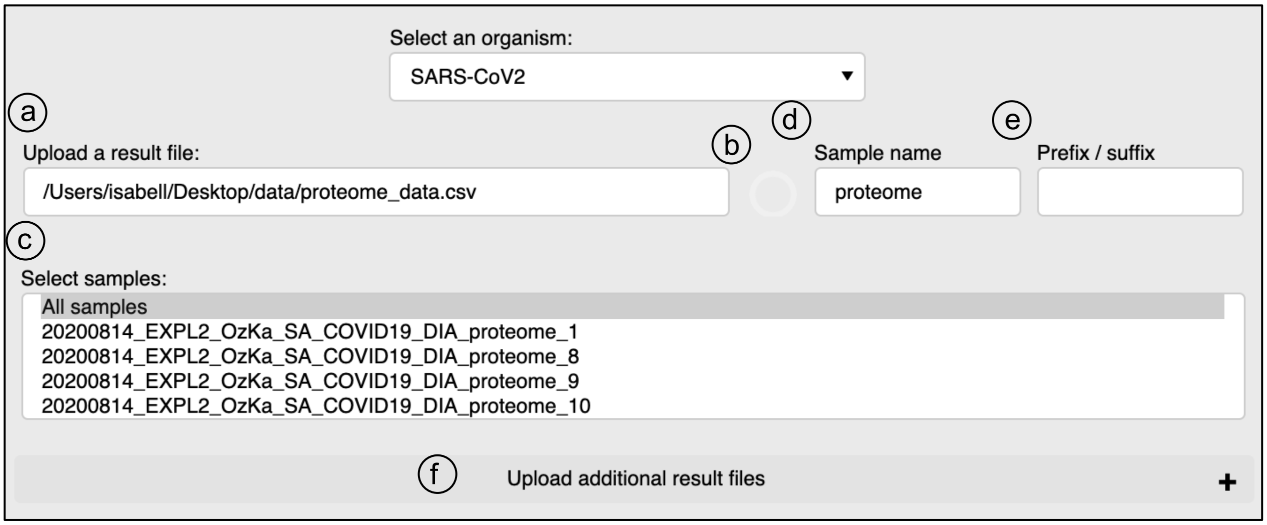

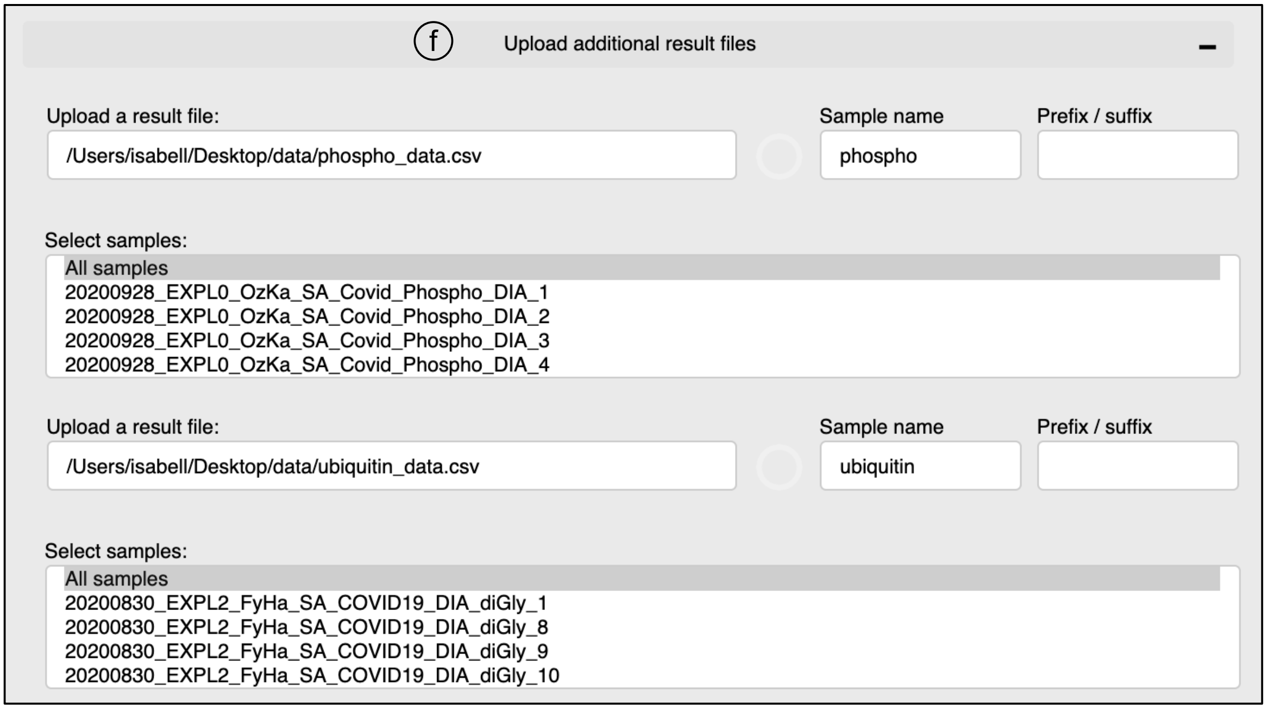


- * If you would like to choose different samples from the same result file, you need to provide the same filepath and select the different samples.
- * To remove the result file that was previously uploaded, just remove its path from the "Upload a result file:" field (2a step).
- * If you cannot upload the selected file, please take a look at the detailed instructions for AlphaPept, MaxQuant, Spectronaut and DIA-NN input formats.

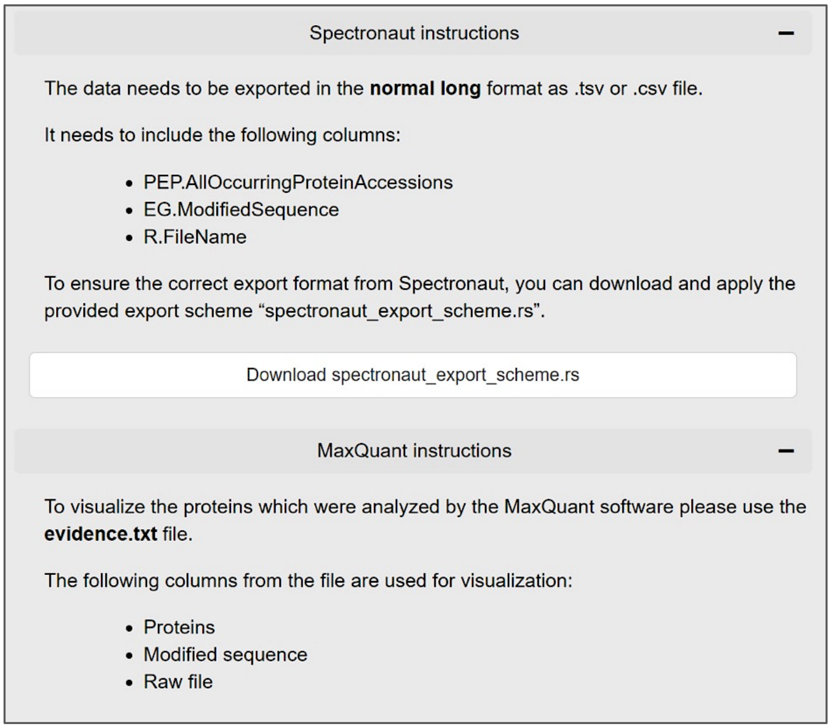

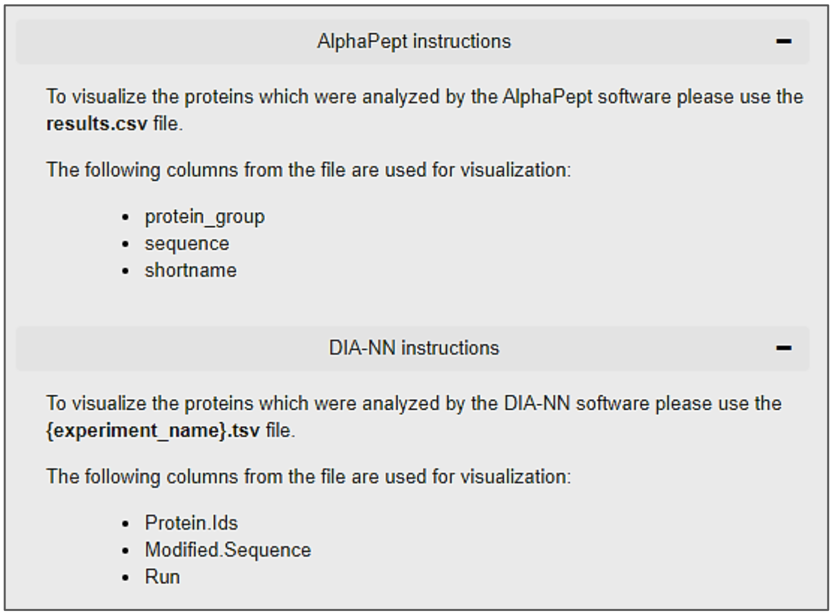


3. Press the "Upload Data" button. The loading process is indicated by a spinner symbol.

4. Select a protein of interest. Per default, you can choose from all UniProt accessions. Click the “Search by a gene name” option to select proteins by their gene name.

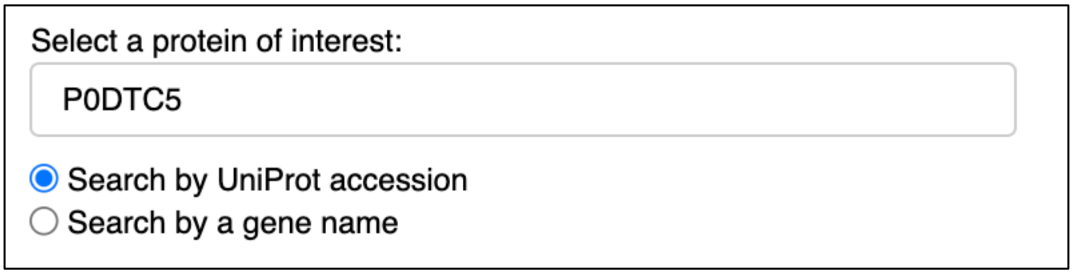


5. (optional) Load a list of pre-selected proteins. This can be a .txt file containing either UniProt accessions or gene names containing one identifier per line. This will reduce the options available in the selection of proteins of interest (step 4). When the list of proteins is loaded, there is an option to generate a pdf report with plots for all pre-defined proteins. The report is intended for fast screening of the data. Note that we generally recommend the interactive visualization and the download of single .svg files as described in step 7a. The report generation process is indicated by a spinner symbol.

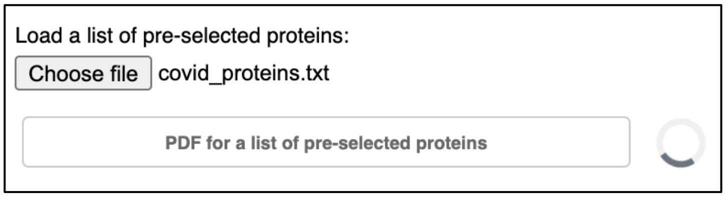

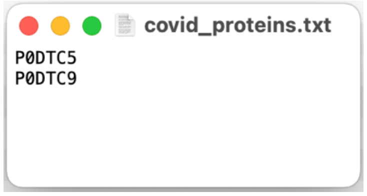

IMPORTANT! Once you uploaded the list of pre-defined proteins, the original list of proteins in the uploaded result file(s) will be filtered. Proteins that are not contained in the list of pre-defined proteins will no longer be available for visualization. If you would like to use the whole list again, repeat from step 2.

6. Select annotation options for the sequence visualization.

    - All sequence annotations from UniProt are available and displayed per default. You can choose a customized set of displayed annotations in the "UniProt annotation" selection.

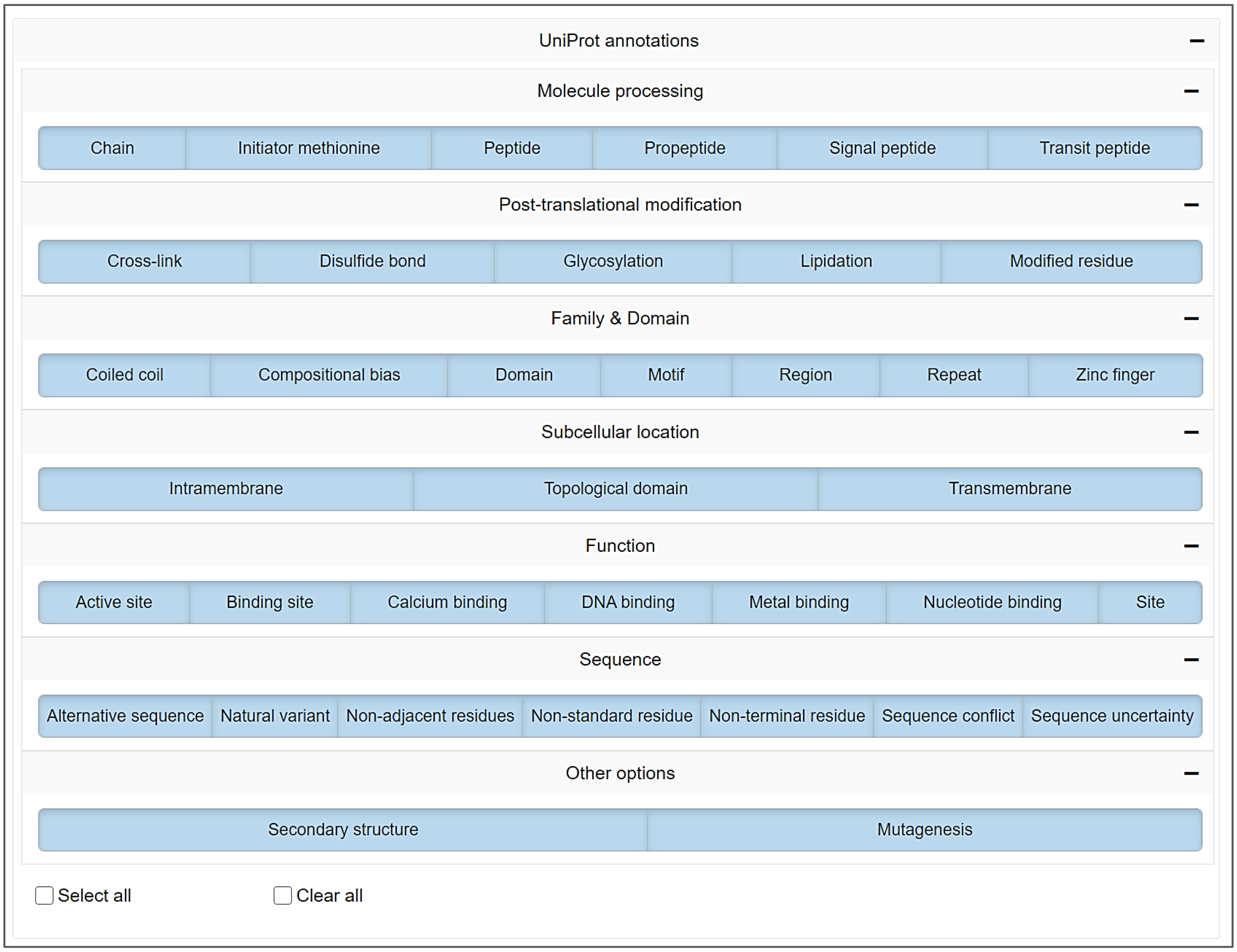


    - All theoretical cleavage sites for the most common proteases can be shown. Trypsin is selected by default. Alternative or additional proteases can be selected in the "Protease cleavage sites" selection.

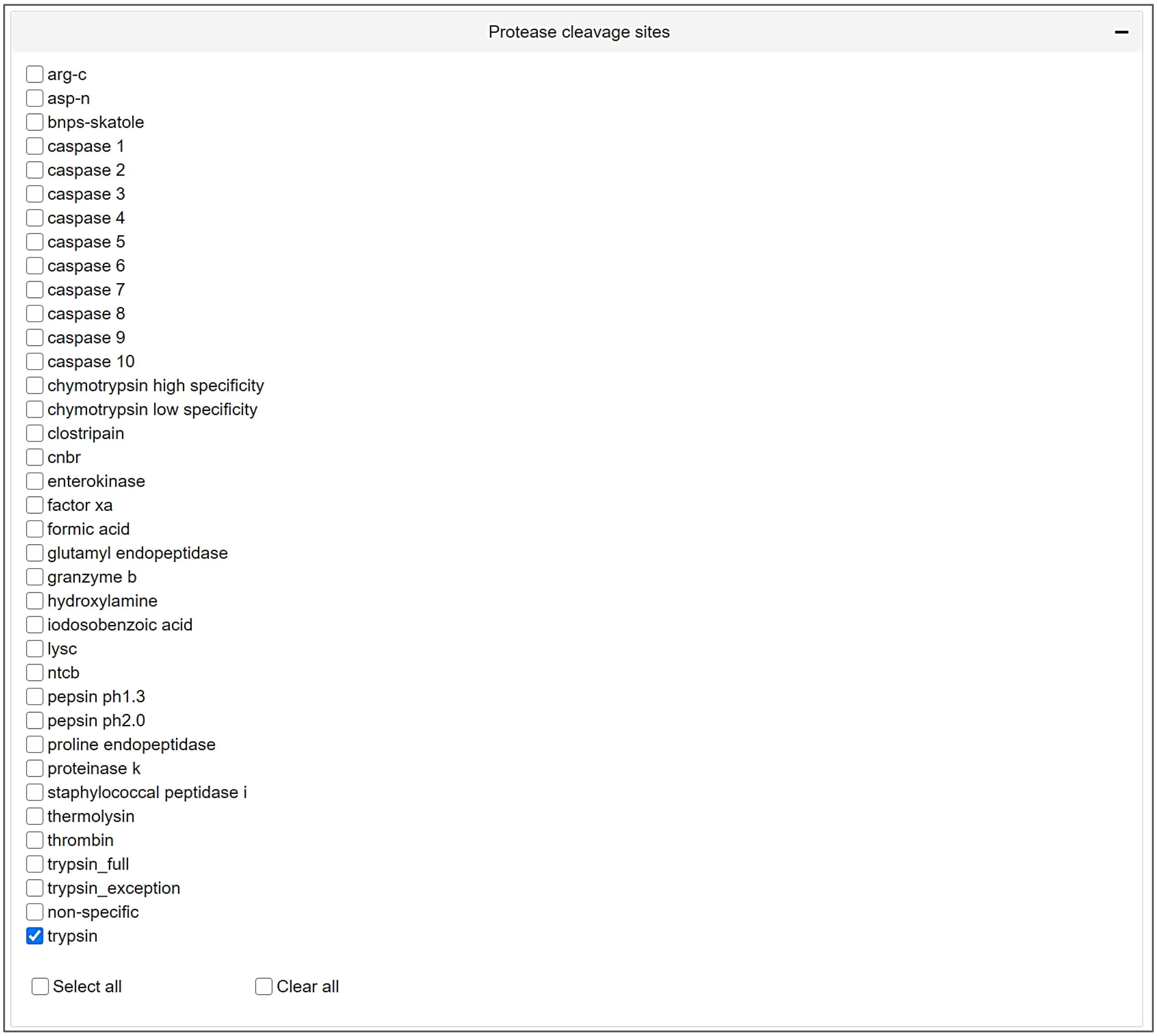


- * You can use "Select all" or "Clear all" checkboxes to speed up the selection process.

7. Press the "Visualize Protein" button. The loading process is indicated by a spinner symbol.

   a) You can use the interactive toolbar to for example zoom in and out or to highlight specific sequence regions. Press the little camera icon to download a high-resolution .svg image of the currently displayed protein and sequence region.

   b) If you hover over the sequence, all annotation information for the current sequence position of the cursor will be displayed.

   c) You can directly visit other websites for further exploration of details on the selected protein of interest. UniProt, PhosphoSitePlus, Protter, PDB an Peptide Atlas are available for direct access.

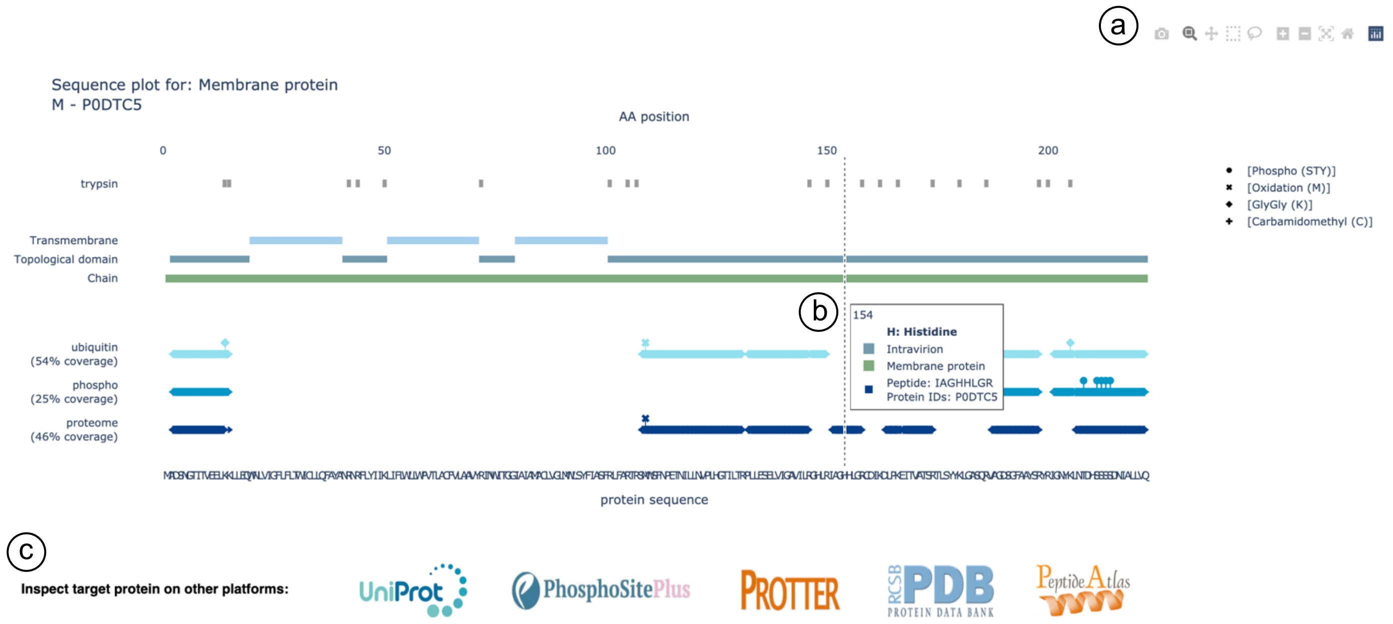


Here we show an example sequence representation of the 'Membrane protein' (P0DTC5) of SARS-CoV-2. The intraviron regions are well covered by the experimental data, whereas none of the observed peptides map to the helical trans-membrane domains. Both phosphorylations ('Phospho (STY)', indicated by circles) and ubiquitinations ('GlyGly (K)', indicated by diamonds) could be identified on the viral protein. This is just one example of how to inspect proteomics data with AlphaMap.

**Now enjoy exploring your own data!**
